# Central nervous system hypomyelination disrupts axonal conduction and behaviour in larval zebrafish

**DOI:** 10.1101/2021.04.20.440476

**Authors:** Megan E Madden, Daumante Suminaite, Elelbin Ortiz, Jason J Early, Sigrid Koudelka, Matthew R Livesey, Isaac H Bianco, Michael Granato, David A Lyons

## Abstract

Myelination is essential for central nervous system (CNS) formation, health and function. As a model organism, larval zebrafish have been extensively employed to investigate the molecular and cellular basis of CNS myelination, due to their genetic tractability and suitability for non-invasive live cell imaging. However, it has not been assessed to what extent CNS myelination affects neural circuit function in zebrafish larvae, prohibiting the integration of molecular and cellular analyses of myelination with concomitant network maturation. To test whether larval zebrafish might serve as a suitable platform with which to study the effects of CNS myelination and its dysregulation on circuit function, we generated zebrafish myelin regulatory factor (*myrf*) mutants with CNS-specific hypomyelination and investigated how this affected their axonal conduction properties and behaviour. We found that *myrf* mutant larvae exhibited increased latency to perform startle responses following defined acoustic stimuli. Furthermore, we found that hypomyelinated animals often selected an impaired response to acoustic stimuli, exhibiting a bias towards reorientation behaviour instead of the stimulus-appropriate startle response. To begin to study how myelination affected the underlying circuitry, we established electrophysiological protocols to assess various conduction properties along single axons. We found that the hypomyelinated *myrf* mutants exhibited reduced action potential conduction velocity and an impaired ability to sustain high frequency action potential firing. This study indicates that larval zebrafish can be used to bridge molecular and cellular investigation of CNS myelination with multiscale assessment of neural circuit function.

## Introduction

Myelination is a well-characterised regulator of axonal health and function. In recent years it has become clear that myelination in the central nervous system (CNS) is dynamically regulated over time, including by neuronal activity, leading to the view that activity-regulated myelination might represent a form of functional plasticity (1). Furthermore, disruption to myelin is observed in numerous diseases of the CNS, and its regulation may represent a viable therapeutic strategy. Indeed, major insights have emerged from studies in multiple systems into the cellular and molecular mechanisms of CNS myelination, its regulation by neuronal activity, and its disruption in disease (2–5). In parallel, an increasing number of studies indicate that the generation of new oligodendrocytes (6–10), and the degree of myelination (11–14), are important for distinct behaviours. However, how dynamic regulation of myelination, or disruption to myelin per se, actually affects the activity of circuits underlying these behaviours remains much less clear. This is partly due to the difficulty in visualising changes to myelination along single axons over time in the mammalian brain while concomitantly assessing their conduction properties, and assessing how alteration to conduction in turn affects neural circuit function.

Zebrafish are well established as a model organism for the study of myelination. The small size and transparency of their larvae, in combination with established transgenic tools, allows the assessment of myelin made by individual oligodendrocytes and along single axons in vivo (15, 16). Combined, with their genetic tractability, many insights into molecular and cellular mechanisms of myelination have come from investigations in this model (17). Despite this progress, it remains unknown how CNS myelination affects the function of individual axons, neural circuits, or the behaviour of larval zebrafish, and thus it is not clear whether integrated multiscale assessments of CNS myelination from molecule through circuit can be performed in this model. However, it is now clear that larval zebrafish exhibit a diverse repertoire of experimentally tractable innate and stereotypical locomotor behaviours (18), many of which are mediated by reticulospinal (RS) neurons – a diverse set of neurons of the midbrain and hindbrain that process multimodal sensory information, and project descending axons to the spinal cord to coordinate specific motor outputs (19, 20). Intriguingly, RS axons are first to be myelinated in the zebrafish CNS and exhibit activity-regulated myelination at larval stages (16, 21, 22), implying that regulation of their myelination might influence circuit function in early larvae. In vivo electrophysiological recordings from subsets of individual RS neurons are feasible (23–25), which in principle permits direct measurement of myelinated axon conduction properties underlying behaviour. However, how disruption to CNS myelination affects the behaviour or axonal conduction properties of larval zebrafish remains to be investigated.

In this study, we set out to investigate whether changes to CNS myelination can be detected in behaviour and in the conduction properties of single axons in zebrafish larvae. To achieve this, we created a myelin gene regulatory factor (*myrf*) mutant line, which exhibits severe CNS hypomyelination. Using this mutant, we demonstrate that both behavioural and electrophysiological consequences of hypomyelination are indeed detectable in the relevant circuitry in vivo, providing proof of principle that integrated analysis is feasible in this model organism, offering a framework for future investigations.

## Results

### Targeting myelin gene regulatory factor to create a larval zebrafish model of CNS-specific hypomyelination

To begin our investigations into the role of CNS myelination in neural circuit function, we sought to establish a larval zebrafish model with CNS-specific hypomyelination. Mammalian studies have identified *myrf* as a transcription factor vital for CNS myelin formation and maintenance (26, 27). Zebrafish possess a single ortholog of *myrf* and, similar to mammals, *myrf* expression in the CNS appears to be restricted to oligodendrocytes (28, 29). We used CRISPR/Cas9 technology to target a guide RNA to exon 2 of the zebrafish *myrf* gene, the first conserved exon across all predicted splice variant isoforms, and in doing so created the *myrf*^*UE70*^ mutant (**Methods** and **Figure 1A**). Morphometric analysis of larval body features of *myrf*^*UE70*^ mutants at larval stages showed them to be indistinguishable from siblings (data not shown), and in contrast to mammalian *myrf* mutants (27), homozygous *myrf*^*UE70*^ mutants remain viable through to adulthood. Adult *myrf*^*UE70*^ mutants exhibited an almost complete absence of myelin basic protein (mbp) mRNA (**Figure 1B**), and transmission electron microscopy (TEM) assessment indicated that had effectively no myelin in the adult spinal cord (**Figure 1C** and **D**). In addition, and unlike larvae, homozygous adult *myrf*^*UE70*^ were grossly identifiable from their siblings by their smaller size. Adult *myrf*^*UE70*^ mutants were also infertile, due to the absence of detectable gonadal tissue in females, confirmed via histopathology, which also revealed evidence of cardiomyopathy (data not shown), findings consistent with proposed roles of *myrf* outside the CNS (30–32).

**Fig. 1.**
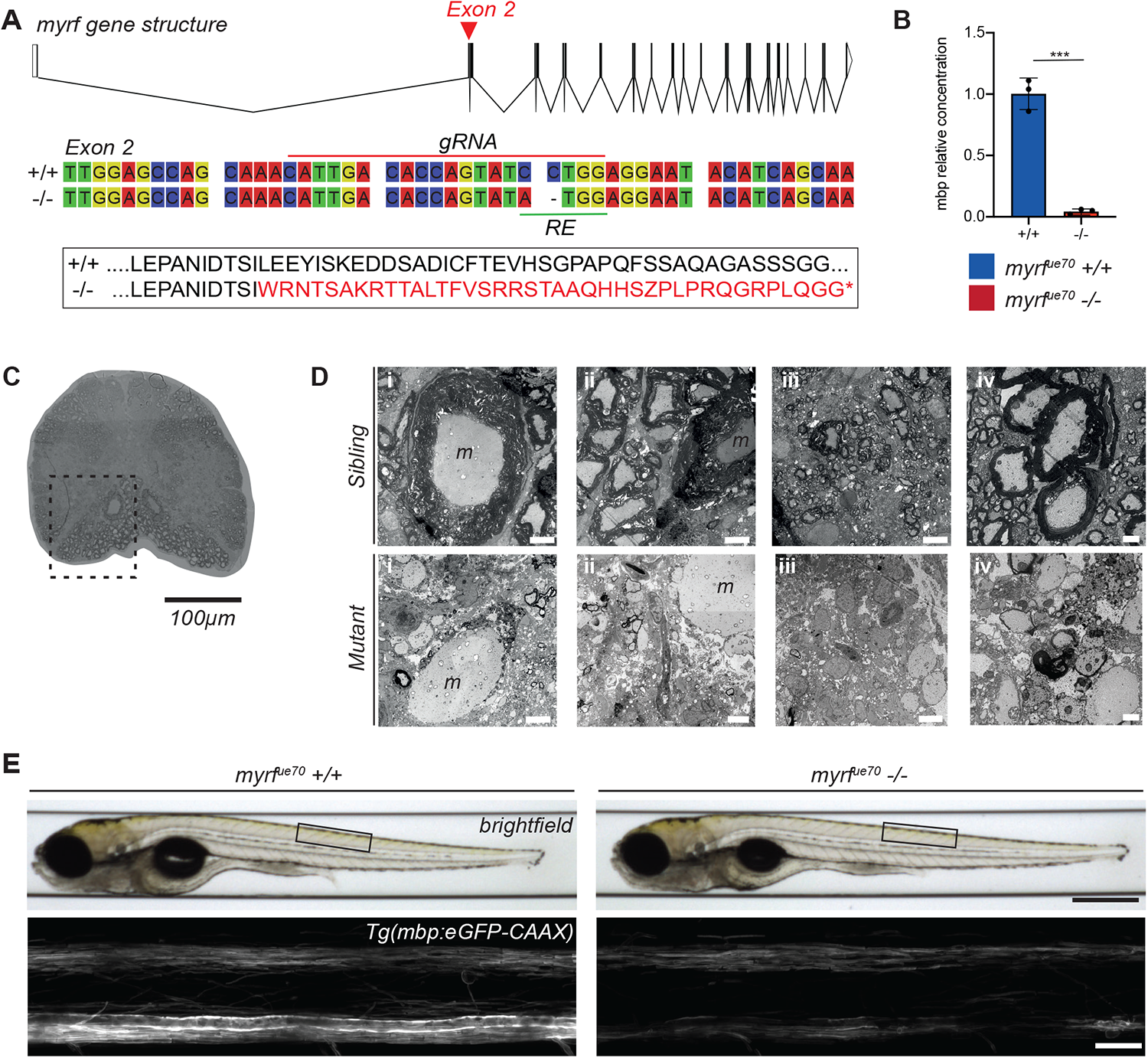
*myrf*^*UE70*^ mutants display a gross reduction in the level of CNS myelination at the adult and larval stages. **A)** Top: *myrf* gene structure composed of 27 exons. Red arrowhead marks the location of the mutation in exon 2. Scale bar equates to 1000bp. Schematic created using Wormweb.org. Middle: Wildtype and mutant nucleotide sequences spanning the mutagenesis site. The guide RNA (gRNA) target site (red line) and restriction enzyme (RE) recognition site (green line) are labelled. Bottom: Amino acid sequence indicating that the *myrf*^*UE70*^ mutation results in shift in the open reading frame leading to downstream coding for a premature stop codon (*). **B)** The relative concentration of mbp mRNA is reduced by 95% in mutants (0.04 ± 0.03 au) compared to wildtypes (1.003 ± 0.13 au, p = 0.0002, unpaired t test, N = 3 adult brains per genotype). **C)** Transverse section of the spinal cord in an adult *myrf*^*UE70*^ sibling showing extensive myelination of ventral spinal cord (dashed box). 20x objective. Scale bar = 100µm. **D)** TEM images of the spinal cord in the region of the ventral spinal tract (outlined in C) in *myrf*^*UE70*^ adult siblings (top) and mutants (bottom). ‘m’ denotes the Mauthner axon. **E)** Top: Brightfield images of *myrf*^*UE70*^ wildtype and mutant larvae at 6dpf. Black box defines the anatomical region imaged across animals. Scale bar = 0.5mm. Bottom: Confocal microscopy images of the spinal cord at 6dpf in *myrf*^*UE70*^ Tg(mbp:eGFP-CAAX) larvae. Scale bar = 20µm. Error bars represent mean ± standard deviation.

Given the potential to study myelination of well-defined circuits at high resolution over time at larval stages when *myrf*^*UE70*^ mutants are morphologically indistinguishable from siblings, we next analysed our transgenic reporter of myelination Tg(mbp:eGFP-CAAX) at 6 days post fertilisation (dpf). This indicated that the gross level of CNS myelination was also reduced in *myrf*^*UE70*^ mutant larvae relative to wildtype siblings (**Figure 1E**). To quantify myelination in larvae, TEM was performed on transverse sections of the spinal cord (CNS) and posterior lateral line nerve of the peripheral nervous system (PNS) at 6 dpf (**Figure 2A-C**). At this timepoint, we observed a 66% reduction in the number of myelinated axons in the spinal cord of *myrf*^*UE70*^ mutants relative to wildtype siblings (35.29 ± 7.83 myelinated axons in wildtypes, 12.00 ± 4.34 myelinated axons in mutants, p ≤ 0.0001, unpaired t-test, **Figure 2D**). In contrast, and demonstrating specificity of hypomyelination to the CNS, similar numbers of myelinated axons were observed in the PNS of mutant and wildtype siblings (7.33 ± 1.53 myelinated axons in wildtypes, 9.00 ± 3.83 myelinated axons in mutants, p = 0.52, unpaired t-test, **Figure 2E**).

**Fig. 2.**
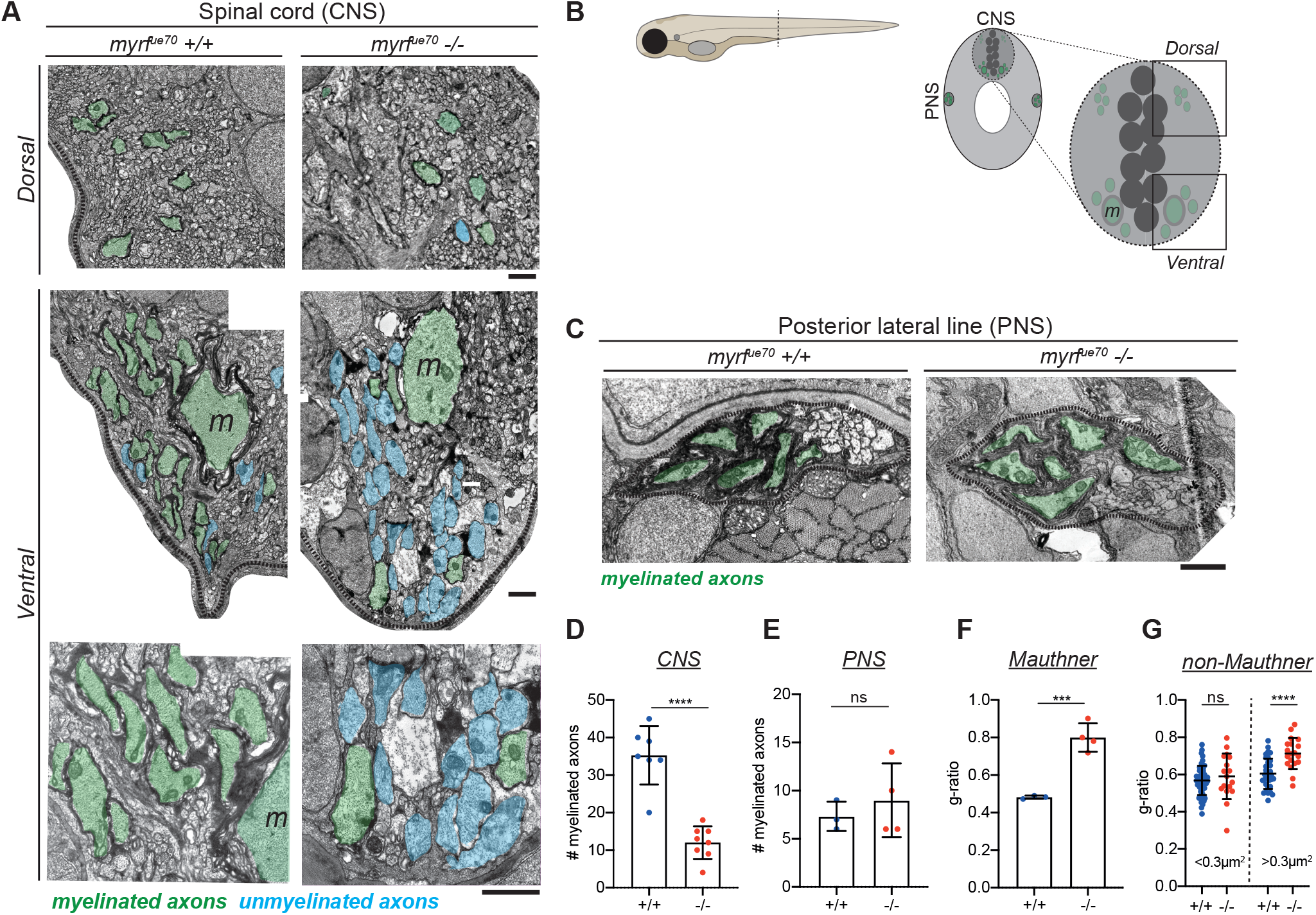
*myrf*^*UE70*^ mutants display CNS-specific hypomyelination. **A)** TEM images of the myelinated tracts in the dorsal and ventral spinal cord, with a high magnification image provided (bottom row). Scale bars = 1µm. **B)** Schematic of the transverse section of a 6dpf larval zebrafish at the level of the urogenital opening. Inset: transverse section of the spinal cord at the same level. Myelinated (green) axons are located in the ventral and dorsal spinal tracts of the spinal cord (CNS) as well as the posterior lateral line (PNS). m = Mauthner axons. **C)** TEM images of the posterior lateral line at 6dpf. Scale bar = 1µm. **D)** The average number of myelinated axons in one hemi-spinal cord is reduced by 66% in mutants (wildtypes: 35.29 ± 7.83 myelinated axons, mutants: 12.00 ± 4.34 myelinated axons, p ≤0.0001, unpaired t-test, N = 7 wildtypes, N = 8 mutants). **E)** The number of myelinated axons in the PNS is similar between genotypes (wildtypes: 7.33 ± 1.53 myelinated axons, mutants: 9.00 ± 3.83 myelinated axons, p = 0.52, unpaired t-test, N = 3 wildtypes, N = 4 mutants). Values represent mean ± standard deviation. **F)** G-ratio of Mauthner axons in wildtype and mutant siblings (wildtypes: 0.48 ± 0.009, mutants: 0.80 ± 0.08, p =0.0009, unpaired t-test). **G)** G-ratios for myelinated axons for small caliber (area <0.3µm^2^) and large caliber (area >0.3µm^2^) myelinated axons. The g-ratio of small caliber axons is similar between groups (wildtypes: 0.57 (0.52 to 0.62), mutants: 0.59 (0.52 to 0.70), p = 0.51, Mann-Whitney test, n = 53 myelinated axons in wildtypes, n = 17 myelinated axons in mutants). The g-ratios for large caliber axons are significantly higher in mutants than wildtype siblings (wildtypes: 0.60 ± 0.08, mutants: 0.71 ± 0.08, p ≤0.0001, unpaired t-test, n = 33 myelinated axons in wildtypes, n = 19 myelinated axons in mutants).

Despite the large number of unmyelinated axons in *myrf*^*UE70*^ mutants, our TEM analyses indicated that some axons remained ensheathed in the larval CNS, including the very large diameter Mauthner axons, the first axons myelinated in the zebrafish CNS (21). Although Mauthner axons were ensheathed in *myrf*^*UE70*^ mutants at 6dpf, they had significantly thinner myelin sheaths compared to wildtype siblings (average g-ratio: 0.48±0.009 in wildtypes, 0.80±0.08 in homozygous mutants, p = 0.0009, unpaired t-test, **Figure 2F**). A similar finding was observed in the other axons that were ensheathed in *myrf*^*UE70*^ mutants at this stage, with greater g-ratio values (denoting thinner myelin) for other large caliber axons in mutants than in wildtype siblings (average g-ratio: 0.60 ± 0.08 wildtypes, 0.71 ± 0.08 mutants, p ≤ 0.0001, unpaired t test, **Figure 2G**). Despite the generally severe hypomyelination phenotype, the presence of some large caliber myelinated axons in zebrafish *myrf*^*UE70*^ mutants at larval stages contrasts with our analysis of adult zebrafish mutants and *myrf* mutant mice which both have essentially a complete absence of CNS myelination (27). This suggests that the full effects of *myrf* knockout may be masked at early stages, either by maternal gene expression or genetic compensatory mechanisms (33).

To examine the cellular basis of CNS hypomyelination in *myrf*^*UE70*^ mutant larvae, we first assessed myelinating oligo-dendrocyte number using the transgenic reporter Tg(mbp:nls-eGFP) (34) (**Figure 3A**). At 6dpf, the timepoint at which TEM was performed, the number of detectable oligodendrocytes was reduced by 21% in *myrf*^*UE70*^ mutants relative to wildtype siblings (p = 0.0002, unpaired t-test, **Figure 3B**). In addition, the fluorescent intensity of *myrf*^*UE70*^ mutant oligo-dendrocyte nuclei was reduced, consistent with reduced mbp expression. Because, the reduction in cell number was not sufficient to explain the reduction in myelin observed using TEM, we assessed the morphology of individual myelinating oligodendrocytes using mosaic cell labelling with the transgenic reporter mbp:mCherry-CAAX (21) (**Figure 3C**). We found that both myelin sheath number (p = 0.02, Mann-Whitney test, **Figure 3D**) and length (p = 0.002, unpaired t test, **Figure 3E**) were reduced in *myrf*^*UE70*^ mutants by 33% and 25% respectively at 6 dpf, with total myelin (sum of sheath lengths) per individual oligodendrocyte reduced by 47% in mutants relative to wildtypes (p ≤ 0.0001, unpaired t test, **Figure 3F**). In addition to being required for the initiation of myelination, previous studies in rodents indicate that *myrf* is also essential for myelin sheath maintenance (35). Having observed that adult *myrf*^*UE70*^ mutants have a much more severe hypomyelination phenotype than larvae (**Figure 1C**), we wanted to assess how the morphology of single oligodendrocytes changed over time. To do so, we imaged single oligodendrocytes at 4dpf and again at 6dpf (**Figure 3G**). We found that between these timepoints mutant oligodendrocytes demonstrated a net shrinkage in myelin sheath length, while wildtype oligodendrocytes showed a net growth (p = 0.009, Mann-Whitney test, **Figure 3H**). Further-more, the number of myelin sheaths that were completely retracted during this timeframe was significantly higher in *myrf*^*UE70*^ mutant oligodendrocytes (p = 0.003, unpaired t test, **Figure 3I**). Also consistent with a failure to maintain healthy myelin sheaths, the number of myelin sheaths exhibiting an abnormal morphology (i.e. incomplete wrapping, abnormal elongation profiles or myelin blebs) was significantly higher in mutant versus wildtype oligodendrocytes at 6dpf (**Figure 3J**).

**Fig. 3.**
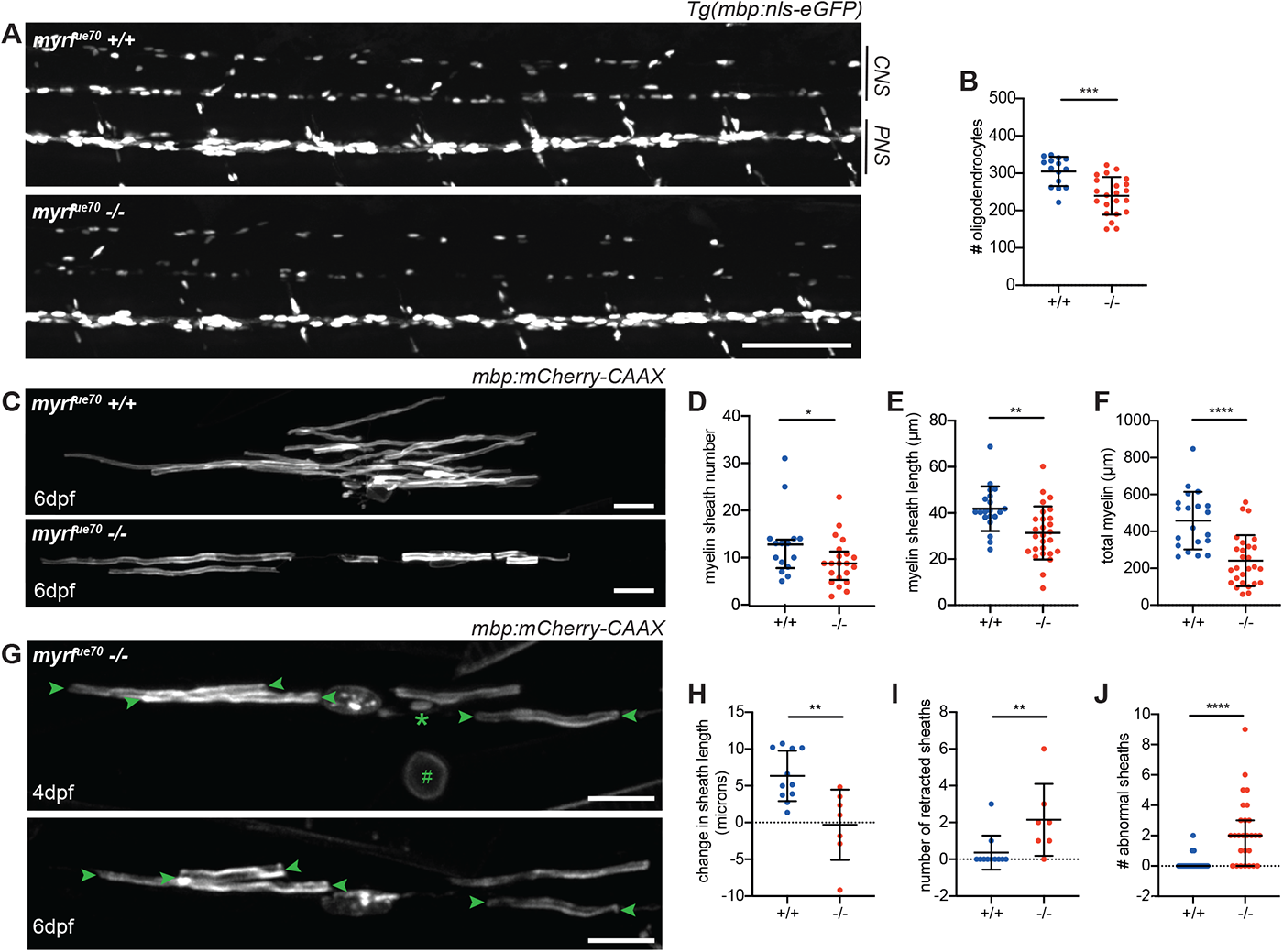
*myrf*^*UE70*^ mutants have fewer oligodendrocytes which produce less myelin and fail to maintain myelin sheaths over time. **A)** Confocal images of the spinal cord at 6dpf in sibling control and *myrf*^*UE70*^ Tg(mbp:nls-eGFP) larvae. Scale bar = 100µm. **B)** Oligodendrocyte numbers in the spinal cord at 6dpf (wildtype: 304.8 ± 39.07, mutants: 239.3 ± 50.48, p= 0.0002, unpaired t-test, N = 15 wildtypes, N = 22 homozygous mutants). Error bars represent mean ± standard deviation. **C)** Representative confocal images of single oligodendrocytes mosaically labelled with mbp:mCherry-CAAX in a wildtype (top) and mutant (bottom) at 6dpf. Scale bar = 15µm. **D)** Average myelin sheath number was reduced in *myrf*^*UE70*^ mutants relative to wildtype siblings at 6dpf (wildtypes: 10.50 (7.00 to 14.00) sheaths per cell, mutants: 7.00 (5.00 to 10.50) sheaths per cell, p = 0.02, Mann-Whitney test). Values and error bars represent median and interquartile range. **E)** Average myelin sheath length was reduced from 41.83 ± 9.68µm in wildtypes to 31.35 ± 11.49µm in mutants at 6dpf (p = 0.002, unpaired t test). Error bars represent mean ± standard deviation. **F)** Total myelin produced per oligodendrocyte was reduced from 458.2 ± 156.4 µm in wildtypes to 241.1 ± 138.6µm in mutants at 6dpf (p ≤0.0001, unpaired t test). Error bars represent mean ± standard deviation. D-F: N = 20 wildtypes, N = 27 mutants. **G)** Confocal images of a single mutant oligodendrocyte labelled with mbp:mCherry-CAAX at 4 and 6dpf. A myelin sheath (*) and myelinated neuronal cell body (#) are observed at 4dpf and subsequently retracted by 6dpf. Arrowheads label myelin sheaths which are observed to shrink between 4 and 6dpf. Scale bar = 15µm. **H)** Myelin sheaths belonging to wildtype oligodendrocytes demonstrated a net growth of 6.24 ± 3.43µm between 4 and 6dpf. While mutants display net shrinkage of myelin sheaths by -0.31 ± 4.79µm (p = 0.003, unpaired t test). Error bars represent mean ± standard deviation. **I)** Between 4 and 6dpf, wildtype oligodendrocytes retracted 0 (0 to 0) myelin sheaths, while mutants retracted 2 (1 to 3) myelin sheaths (p = 0.009, Mann-Whitney test). Error bars represent median and interquartile range. **J)** Number of abnormal myelin sheaths at 6dpf (wildtypes: 0.00 (0.00-0.00) abnormal sheaths; mutants: 2 (0.00-3.00) abnormal sheaths, p ≤0.001, Mann-Whitney test). Error bars represent median and interquartile range. H – I: N = 11 wildtypes, N = 7 mutants.

In summary, disrupting *myrf* leads to a CNS-specific hypomyelination phenotype in larval zebrafish, caused by a reduction in the number of oligodendrocytes, with those that remain having fewer and shorter sheaths. The majority of sheaths that are made are thinner, and, based on our documentation of almost complete absence of myelin in adults, not maintained long-term. Therefore, the phenotype in the *myrf*^*UE70*^ mutant fulfilled our aim to generate a CNS-specific model of hypomyelination to study the effects on neural circuit function at larval stages.

### *myrf*^*UE70*^ mutants exhibit an increase in the latency to perform startle responses and an impaired behavioural choice in response to a defined auditory stimulus

Given that many larval zebrafish sensorimotor behaviours are mediated by RS neurons, whose axons are myelinated early and exhibit activity-regulated myelination (16), we hypothesised that *myrf*^*UE70*^ mutants would display detectable differences in the performance of RS-mediated behaviours. To test this, we chose to first examine acoustic-startle behaviour, for which the underlying circuit is relatively well described (36). Briefly, a high-intensity acoustic stimulus activates the auditory (VIIIth) nerve, which courses into the hindbrain to synapse onto the Mauthner cell at its lateral dendrite. Once the threshold potential is exceeded, an action potential is elicited and rapidly propagated along the Mauthner axon, which crosses into, and extends along, the contralateral tract of the spinal cord. Along its length, collateral branches make synapses with interneurons and primary motor neurons that coordinate motor output. Activation of a Mauthner axon results in a stereotypical, high-velocity ‘c-bend’ away from the stimulus, followed by a fast burst swim (37) (**Figure 4A**). The latency to perform such a response is defined as the time taken from stimulus presentation to the onset of a c-bend (**Figure 4J**). Given that myelin increases conduction velocity along a single axon (38), we made the prediction that the latency to execute the motor responses following an acoustic stimulus would be delayed in *myrf*^*UE70*^ mutants.

**Fig. 4.**
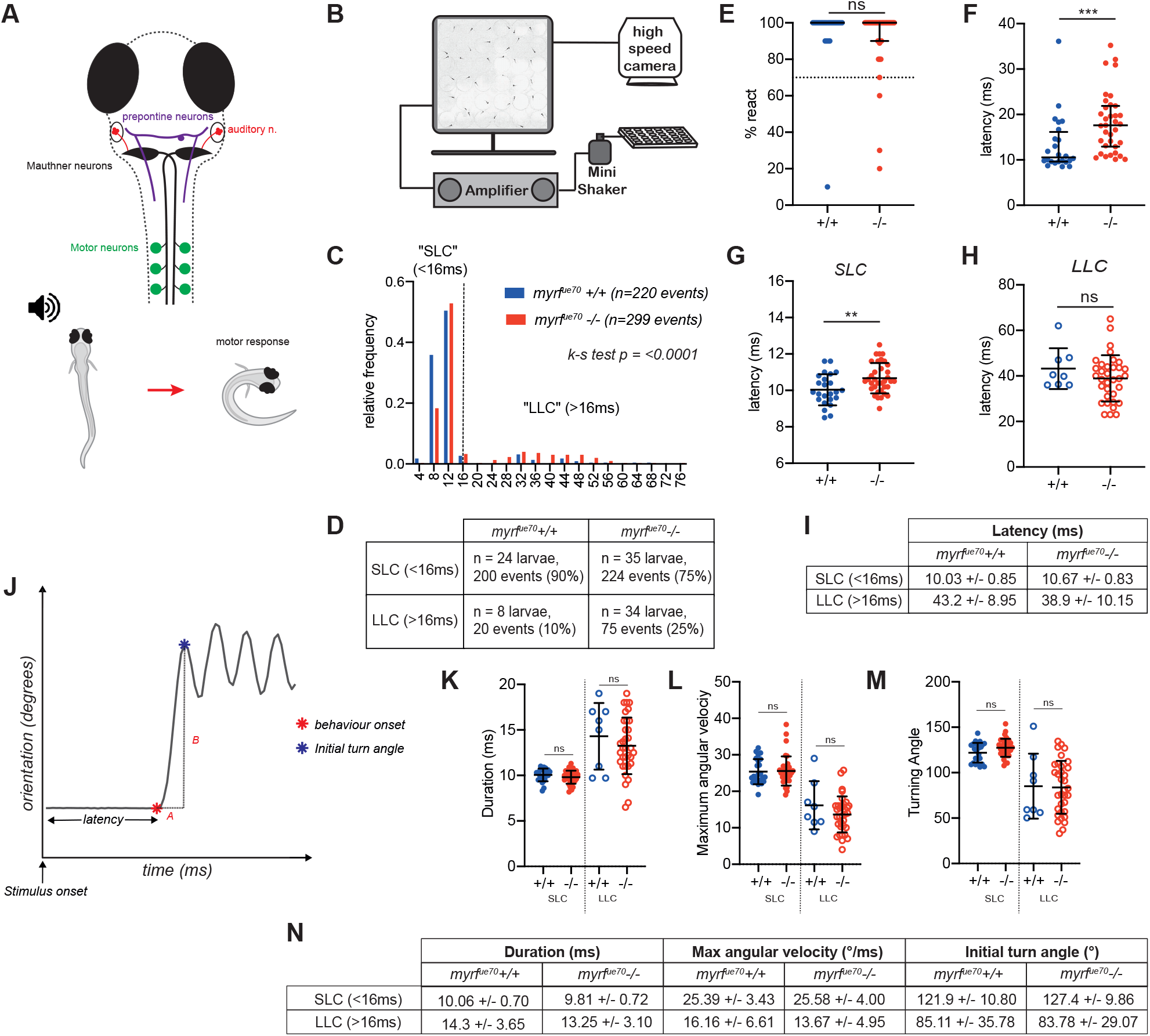
*myrf*^*UE70*^ mutants exhibit increased latency to perform startle responses, and a tendency to perform avoidance behaviour, in response to defined acoustic stimuli. **A)** Overview of the neuronal circuitry involved in motor response to auditory stimuli. **Startle response (SLC):** sensory input from the ear, via the auditory nerve (red) is received at the lateral dendrite of the Mauthner cell body (black). The axon of the Mauthner cell crosses into the contralateral aspect of the spinal cord where it extends along the ventral tract to recruit motor neurons directly along the length of the larvae. Recruitment of motor neurons allows muscle contraction on the side of the body contralateral to the stimulus, allowing a rapid, high-velocity c-bend (motor response) away from the stimulus. **Avoidance behaviour (LLC):** sensory input is detected by prepontine neurons (purple) in the hindbrain, which recruit ipsilateral motor neurons indirectly, resulting in a low-velocity, longer latency, c-bend away from the stimulus. **B)** Schematic of the behavioural rig. **C)** Relative frequency histogram displaying the distribution of latencies for behavioural responses in response to acoustic stimuli in wildtype and mutant larvae (N = 24 wildtype larvae, n = 220 events; N = 35 mutant larvae, n = 299 events; Kolmogorov-Smirnov test, p ≤0.0001). **D)** Number and proportion of events (SLC vs LLC) per genotype. **E)** React rate per fish (median react rate = 100% in both wildtypes and mutants, p = 0.24, Mann-Whitney test, N = 25 wildtype larvae, N = 38 mutant larvae). Larvae are excluded from subsequent analysis if they exhibit a react rate <70%. **F)** Average latency values per fish (wildtype: 10.55ms (9.6-16.15ms), mutants: 17.6ms (12.9-21.88ms), p = 0.003, Mann-Whitney test). **G)** Average latency of short latency c-starts (<16ms) (wildtypes: 10.03 ± 0.85ms, mutants: 10.67 +/-0.83ms, p = 0.006, unpaired t test). **H)** Average latency of long latency c-starts (>16ms) (wildtypes: 43.20 ± 8.95ms; mutants: 38.91 ± 10.15ms, p = 0.28, unpaired t test). **I)** Mean and standard deviations values for SLC and LLC responses per genotype. **J-M analysis of c-bend kinematics: J)** example trace of orientation over time during a behavioural response to an acoustic stimulus. C-bend kinematics are calculated from individual traces for each response per fish. Latency is the time from stimulus onset to behavioural onset (red star). C-bend duration (A) is time from behaviour onset to initial turn angle (blue star). Maximum angular velocity is defined as the change in orientation over time (B/A). Turning angle equates to the initial turn angle. **K)** Initial turn duration (SLC: wildtypes: 10.06 ± 0.70ms, mutants: 9.81 ± 0.72ms, p = 0.20, unpaired t-test; LLC: wildtypes: 14.30 ± 3.65ms, mutants: 13.25 ± 3.10, p = 0.42, unpaired t test). **L)** Maximum angular velocity (SLC: wildtypes: 24 (22.78-28.68) °/ms, mutants: 25 (23.10-26.60) °/ms, p = 0.73, Mann-Whitney test; LLC: wildtypes: 16.16 ± 6.61°, mutants: 13.67 ± 4.95°, p = 0.24, unpaired t-test). **M)** Initial turn angle (SLC: wildtypes: 121.9 ± 10.80, mutants: 127.4 ± 9.86, p = 0.051, unpaired t test; LLC: wildtypes: 85.11 ± 35.78, mutants: 83.78 ± 29.07, p = 0.91, unpaired t test). **N)** Descriptive statistics (mean ± standard deviation) for c-bend kinematics. For **Figures E-G** and **K-M**, N = 23 wildtypes, N = 35 mutant larvae. For **Figures D & E** values represent median and interquartile range, for **Figures F-K**, values represent mean ± standard deviation.

Motor behaviour was assessed using an established high-throughput assay (39). *myrf*^*UE70*^ larvae were arrayed into individual wells of a 6×6 custom made plate attached to an amplifier delivering a series of acoustic stimuli at 20 second intervals (**Figure 4B**). Using a high-speed (1000Hz) camera, behavioural responses were recorded and subsequently analysed using FLOTE software (39, 40). Overall, the frequency of responses to acoustic stimuli was similar between groups (**Figure 4E**). However, on average, *myrf*^*UE70*^ mutants exhibited a 66% increase in their average latency to elicit a response compared to wildtype siblings (wildtypes: 10.55ms (9.6-16.16ms), mutants: 17.60ms (12.90-21.88ms), p = 0.003, Mann-Whitney Test, **Figure 4F**).

Interestingly, larval behavioural responses to acoustic stimuli can be modulated across variable stimulus properties, exhibiting decision-making capabilities of the underlying circuitry (39, 41). For example, in larval zebrafish, while high intensity threatening stimuli induce the short-latency c-bend startle response, also known as the ‘short latency c-start’ (SLC), lower stimulus intensities induce a distinct longer latency reorientation-like behaviour, initially defined as a ‘long latency c-start’ (LLC). These kinematically and behaviourally distinct responses are executed by activity in partially overlapping circuitry, with the crucial difference that SLCs are driven by recruitment of Mauthner neurons, while LLCs appear to be driven by alternative pathways e.g. prepontine neurons (39, 42) (**Figure 4A**). Given that hypomyelination in *myrf*^*UE70*^ is widespread within the CNS, we anticipated that the large overall increase in latency to respond to acoustic stimuli might be due to significant delays in the performance of both SLC and LLC responses. However, when data was segmented into SLCs or LLCs, the latency to perform an SLC was increased by 6.4% (10.03 ± 0.85ms in wild-types, 10.67 ± 0.83ms in mutants, p = 0.006, unpaired t test, **Figure 4G** and **I**) and that the latency to perform LLCs remained unaffected (**Figure 4H** and **I**), begging the question as to what caused the much larger overall increase in latencies to respond to acoustic stimuli.

We reasoned that if the latency to perform SLCs was only affected to a small degree and to perform individual LLCs not at all, the overall large increase in latency to perform all responses might be due to a biased selection of the longer latency LLCs over the much shorter latency SLCs. Indeed, when we compared their relative frequency, we saw that LLCs represented a significantly increased proportion of behavioural responses in *myrf*^*UE70*^ mutants relative to wild-types (SLC:LLC ratio: 10:1 in wildtypes, 2.9:1 in mutants, p ≤0.0001, Kolmogorov-Smirnov test, **Figure 4C** and **D**). To ensure that this apparent bias in behavioural selection was not due to SLCs simply being so slow as to be detected as LLCs, we analysed additional kinematic parameters (**Figure 4J-N**), which are specific to each type of response (39). No differences were found in the duration, maximum angular velocity or initial turning angle of SLCs or LLCs between wildtype and mutant larvae (**Figure 4K-M**), consistent with the conclusion that the increased frequency of LLCs represents true LLC events, rather than delayed and inappropriately classified SLCs. In summary, we have shown that *myrf*^*UE70*^ mutants exhibited delayed latency to perform Mauthner-mediated startle responses (SLCs), and an unexpected bias towards performing Mauthner-independent reorientation behaviours (LLCs) in response to the same acoustic stimuli. This shows that hypomyelination in the larval zebrafish can be detected in overt changes to behaviour and highlights the complexity of how dysregulation of myelination impacts circuit function, even when executing relatively simple sensorimotor transformations.

### Action potential conduction is impaired along the Mauthner axon in *myrf*^*UE70*^ mutants

In order to investigate how myelination affects conduction along larval zebrafish axons, we set out to establish an electrophysiological platform that would allow us to measure and compare multiple aspects of axonal conduction in vivo. We focussed our analysis on the Mauthner neuron and axon, due to its characteristic morphology and anatomical location and given its established role in mediating the SLC. To begin with, we performed whole-cell current-clamp recordings of the Mauthner neuron cell body while stimulating its axon in the spinal cord with an extracellular electrode (**Figure 5A**). We first tested whether loss of *myrf* function affected intrinsic properties of the Mauthner neuron, by assessing its resting membrane potential: we found that this remained stable in mutants (siblings: -70.82mV ± 2.76mV, mutants: -70.68 ± 1.25mV, p = 0.9077, unpaired t-test, **Figure 5B**). Our experimental configuration allowed us to record antidromic action potentials propagating along the Mauthner axon. Therefore, we next assessed whether the shape of action potentials along Mauthner axon was disrupted by hypomyelination, given the possibility that this may influence the ability to evoke responses in target neurons. To do so we measured the width of the action potential at its half-height (action potential half-width) at 6dpf and found this to be similar in control and *myrf*^*UE70*^ mutant animals (siblings: 0.64 ± 0.09ms, *myrf*^*UE70*^ mutants: 0.60 ± 0.06ms, p = 0.2610, unpaired t-test, **Figure 5C** and **D**). These data indicate that the degree of hypomyelination along Mauthner axons in *myrf*^*UE70*^ mutants at these stages does not affect the Mauthner resting membrane potential or greatly affect the shape of the action potentials conducted.

**Fig. 5.**
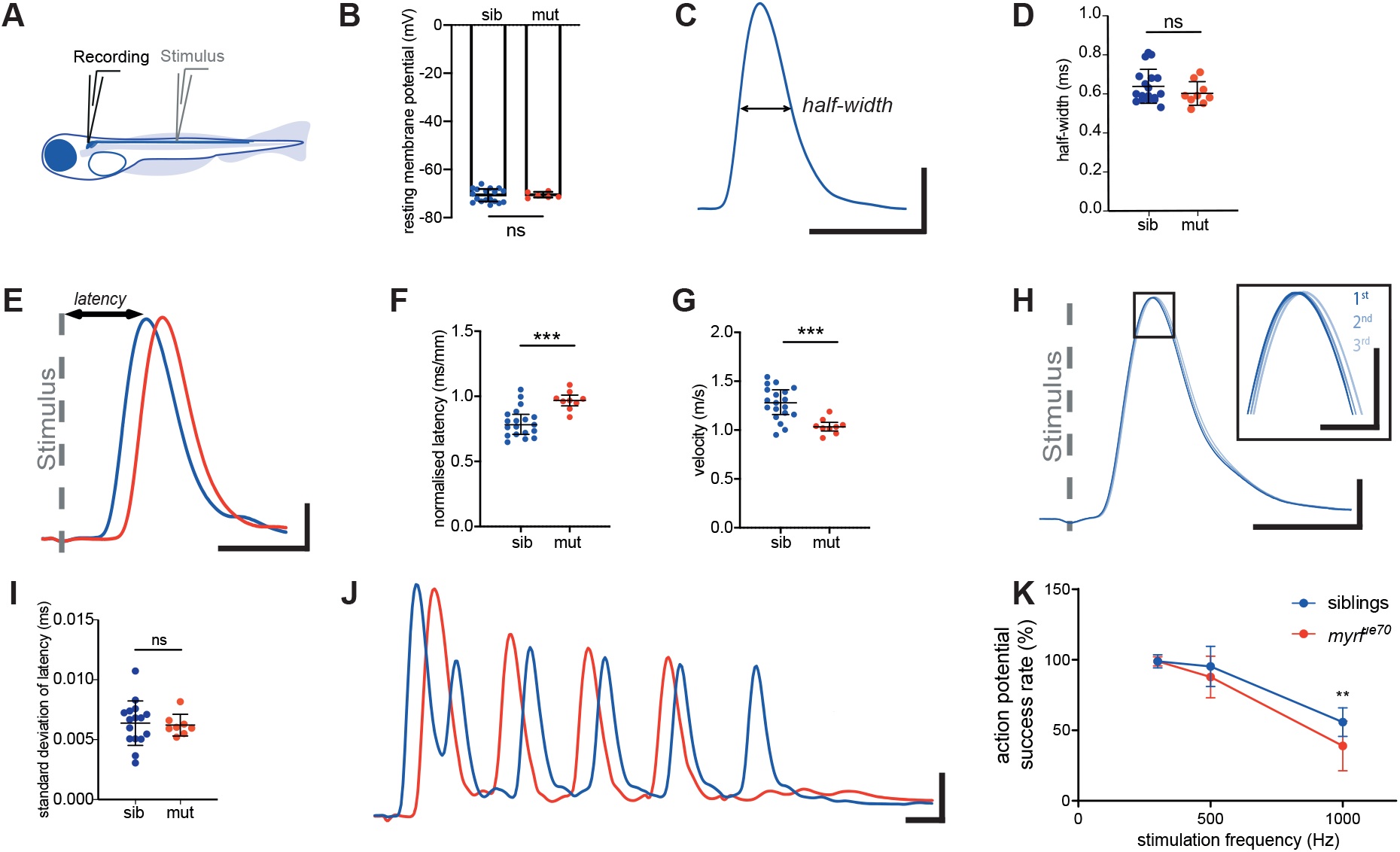
Whole cell current-clamp recordings from Mauthner cells demonstrate slower conduction velocity times and abnormal spiking profiles in *myrf*^*UE70*^ mutants. **A)** Electrophysiological preparation for recording from Mauthner neuron in a whole-cell current clamp configuration while stimulating with an extracellular monopolar field electrode midway through the spinal cord. **B)** Resting membrane potential is unchanged in (siblings (n = 18 cells): -70.82 ± 2.76mV, mutants (n = 6 cells): -70.68 ± 1.25mV, p = 0.9077 at 6 dpf. **C)** Sample trace of an action potential recorded at 6 dpf in a wildtype fish illustrating the measurement of half-width. Half-width is described as width of action potential (ms) and its half height. **D)** Half-width of action potential is unchanged (siblings (n = 18 cells): 0.64 ± 0.09ms, mutants (n = 9 cells): 0.60 ± 0.06ms, p = 0.2610 at 6 dpf). **E)** An example of current – clamp recording from Mauthner neuron in a 6dpf wildtype and mutant following field stimulation (stimulus artefact is indicated by a grey dashed line). Latency is described as time from the onset of stimulus artefact to the peak of action potential. **F)** Normalised action potential latency is increased in mutants at 6dpf (siblings (n = 19 cells): 0.80 ± 0.11ms/mm, mutants (n= 9 cells): 0.97 ± 0.07ms/mm, p = 0.0003 at 6 dpf). **G)** Conduction velocity of Mauthner action potentials are significantly decreased in mutant larvae (siblings (n = 19 cells): 1.27 ± 0.17m/s, mutants (n = 9 cells): 1.04 ± 0.08m/s, p = 0.0005 at 6 dpf). **H)** Sample traces of three subsequent action potentials recorded from the same wildtype Mauthner cell at 6 dpf superimposed and aligned to the peak of stimulus artefact. The area outlined by the rectangle is magnified in the inset and demonstrates imprecise arrival of action potentials. **I)** Precision of action potential arrival is comparable in siblings and mutants (siblings (n = 16 cells): 0.0064 ± 0.0019ms, mutants (n = 8 cells): 0.0062 ± 0.0009ms, p = 0.8166 at 6 dpf). **J)** Sample trace of a train of action potentials fired following 10 stimuli at 1000 Hz at 6 dpf in a *myrf*^*UE70*^ mutant and sibling. **K)** Mauthner neurons in mutant larvae do not sustain prolonged action potential trains of high frequency stimulation (siblings (n = 19 cells): 55.79 ± 10.17% mutants (n = 9 cells): 38.89 ± 17.64% at 6 dpf, p = 0.0014 at 6 dpf). B, D, F, G, I Unpaired t-test. Error bars represent mean ± standard deviation K) Two-way ANOVA. Error bars represent mean ± standard deviation. Scale bars = 10mV and 1ms, except for the inset in H (5 mV and 200µs).

Given the well-defined role for myelin in speeding-up action potential conduction, and the evidence of an increased latency in performing the Mauthner-dependent SLC response, we next measured the latency of action potential conduction along the Mauthner axon in controls and *myrf*^*UE70*^ mutants. This analysis showed that the normalised latency of action potentials was significantly increased in *myrf*^*UE70*^ mutants when compared to siblings (siblings: 0.80 ± 0.11ms/mm, mutants: 0.97 ± 0.07ms/mm, p = 0.0003, **Figure 5F**) resulting in an 18% reduction in conduction velocity (siblings: 1.27 ± 0.17m/s, mutants: 1.04 ± 0.08m/s, p 0.0005, **Figure 5G**). This reduction in conduction velocity supports our finding of a delayed execution of SLCs in *myrf*^*UE70*^ mutants. We next assessed whether the precision of action potential propagation might be impaired due to hypomyelination, which might interfere with synaptic signalling in the circuit. To do so, we measured the ‘jitter’, or imprecision, in the timing of action potential arrival following stimulation, as the standard deviation of 30 action potential peak times aligned to the stimulus artefact (**Figure 5H**). No differences were observed in the precision of action potential arrival in *myrf*^*UE70*^ mutants at 6 dpf (siblings: 0.006 ± 0.002ms, mutants: 0.006 ± 0.0009ms, p = 0.8166, unpaired t-test, **Figure 5I**). These data suggest that hypomyelination leads to slower, but precise, action potential propagation.

Given that the action potentials conducted along Mauthner axons in *myrf* mutants are likely to be sufficient to trigger downstream motor output, albeit with a longer delay, we next asked whether the hypomyelination of Mauthner axon might lead to an increased failure to reliably propagate action potentials. Therefore, we implemented a strategy to robustly test the ability of the myelinated axon to faithfully transmit action potentials. With our preparation, we observed that the Mauthner cell could spontaneously fire short trains of action potentials (1-5) at high frequency (100Hz). We therefore established a high-frequency stimulation paradigm whereby 10 stimuli were delivered in rapid succession via the stimulating electrode and the number of action potentials fired by the cell counted as ‘successes,’ allowing us to assess action potential success rate (**Figure 5J**). Given that myelination reduces axonal current leakage, we predicted that our high frequency stimulation protocol may reveal failed action potential propagation. When we analysed the success rate of action potential firing, we found that this was indistinguishable between siblings and mutants at 300Hz, insignificantly different at 500Hz, but significantly impaired at 1000Hz stimulation, where we found that Mauthner cells from siblings fired at a mean 55.79% success rate, while the mutants were significantly less successful (siblings: 55.79 ± 10.17, mutants: 38.89 ± 17.64, p = 0.0014, two-way ANOVA, **Figure 5K**). This assay indicates that hypomyelination impairs the ability of axons to propagate action potentials faithfully, which could potentially account for the behavioural shift away from Mauthner-mediated responses to auditory stimuli.

In conclusion, we have established an electrophysiology platform that allows direct measurement of single cell (i.e. Mauthner) conduction properties in vivo. In doing so, we have demonstrated that hypomyelination of the Mauthner axon leads to slowed conduction velocity, and with a high frequency stimulation paradigm we reveal a loss of fidelity of action potential propagation along the hypomyelinated Mauthner axon.

## Discussion

We have demonstrated that CNS hypomyelination leads to behavioural alterations and impaired conduction along axons in larval zebrafish. We found that *myrf*^*UE70*^ mutant zebrafish larvae with CNS-specific hypomyelination exhibited an increased latency to execute the stereotypical rapid acoustic startle responses (SLCs) and were also biased towards performing longer latency reorientation behaviours (LLCs) in response to startle-inducing acoustic stimuli. The fact that our analysis revealed phenotypes in both the speed of executing a specific behaviour and in the selection of the correct behavioural response to a sensory stimulus indicates the complex roles that myelination plays in regulating circuit function, providing encouragement that studying additional behaviours will provide further entry-points into studying how alterations to myelination affect the function of other neural circuits. Indeed, there are now a large number of behavioural paradigms that allow analysis of larval zebrafish circuit function, from various sensorimotor transformations (43–45), behaviours regulated by sensory experience over time (39, 46) and those driven by inter-individual interactions, such as sociability (47, 48).

In addition to studying behaviour, we established electrophysiological protocols to assess the conduction properties of single neurons and axons, focusing on the Mauthner neuron due to the ease of its identification and its involvement in the acoustic startle response. We found that conduction along the hypomyelinated Mauthner axon was reduced, and that Mauthner axons in *myrf*^*UE70*^ mutants exhibited an increased failure to propagate action potentials in response to high-frequency stimulation. In our antidromic preparation, we cannot rule out the possibility that the reduced success rate in action potential propagation upon high frequency stimulation was influenced by impaired generation of action potentials. Establishing methods to record orthodromic action potentials remains an important challenge for the future. In addition, electrophysiological analyses of other distinctly identifiable neuronal subtypes and axons, as well as paired recordings of neurons known to communicate within circuits will provide further insight into how myelination and its dysregulation affects the conduction properties of axons and synaptic communication between neurons. Although our study documented behavioural alteration and disruption to conduction in larvae with CNS-specific hypomyelination, many challenges remain in integrating our understanding of circuit function across these scales. However, we believe that the zebrafish represents a model in which a multi-scale functional analyses of myelination on neural circuit function is feasible.

The larval zebrafish brain is relatively simple compared to that of most mammalian models, but nonetheless, has approximately one hundred thousand neurons by 6dpf, and of those, only a relatively small proportion (on the order of a few hundred neurons) have myelinated axons at this stage (49). Furthermore, although the myelination of those axons is generally very stereotyped, it remains adaptable, with some axons exhibiting myelination responsive to neuronal activity. With the aim of studying myelination from the perspective of neural circuits, we previously developed tools to study patterns of myelination along single axons in vivo (16). These tools, together with increasing availability of neuron-specific drivers coupled with circuit maps of the larval fish brain provide a great opportunity to map myelination at single cell resolution across an entire vertebrate CNS, and to do so over time. Even with myelination patterns mapped, a corresponding challenge will be to manipulate myelin from the point of view of specific neurons/axons and circuits. For example, in this study, although we detected millisecond-scale delays in both the latency of the startle response and conduction along the Mauthner axon, it remains unclear whether the longer latency to execute the startle response is simply due to hypomyelination of Mauthner axons, or whether it could be due to conduction delays elsewhere in the circuit, or indeed to more complex integrative functions that affect timing across the circuit. Therefore, it will be important to develop methods to regulate myelination in a neuron/axon and circuit-specific manner. One possibility is to selectively ablate oligodendrocytes in specific circuits. Although oligodendrocyte ablation can be carried out at single cell resolution in zebrafish larvae (15), it leads to inflammatory reactions by cells such as microglia (34), which may be relevant to disease contexts, but would confound the disentangling of the role of myelin per se in healthy circuits. Therefore, an additional approach might be to express cell surface proteins that inhibit myelination (50) along the axons of specific neuronal cell types (51– 53), selectively preventing their myelination. Furthermore, as signals and receptors that influence adaptive myelination are identified, yet more strategies to influence myelination in localised manners may emerge.

In addition to needing more refined methods to map and manipulate myelination of specific circuits, additional tools to assess function across scales from single axon to behaving animal will be required. Complementing behavioural and electrophysiological approaches, optical manipulation and imaging approaches have already proven hugely powerful in the study of larval zebrafish brain function. For example, two-photon and light-sheet microscopy-based imaging studies allow the analysis of neuronal activity from single cells (54) through to sampling the activity of effectively all neurons the entire larval zebrafish brain, at multiple volumes per second with sub-cellular resolution (55–57). In fact, sophisticated imaging platforms that allow monitoring of neuronal activity in the brain during the execution of behaviours have been developed, including during acoustic stimulus-driven responses (41, 58). Furthermore, the coordinated activity of ensembles of neurons have been investigated in the larval brain, which provides an opportunity to investigate how potentially even subtle alterations to myelination in development, health or disease might influence relatively high-order network activity (59–62). To date, most optical analyses of neuronal activity in zebrafish have been carried out using genetically encoded Ca^2+^ reporters, but the limited temporal kinetics of even the fastest Ca^2+^ reporters may preclude the analysis of millisecond-scale changes to conduction properties, which our data indicate can be expected with disruption to larval myelination. However, ongoing development and refinement of voltage indicators appear to exhibit photodynamic properties with the sensitivity to detect functional changes to conduction and synaptic properties at the appropriate temporal resolution, including in larval zebrafish (54). In summary, our study presents larval zebrafish as a viable model to study myelination across scales from molecular and cellular analyses of how myelin organises and supports axons through to functional assessments of conduction, synaptic communication, network function and behaviour over time.

## Acknowledgements

This work was supported by Wellcome Trust Senior Research Fellowships (102836/Z/13/Z and 214244/Z/18/Z) to DAL, a Wellcome Trust Edinburgh Clinical Academic Track PhD studentship (205042/Z/16/Z) to MEM, National Institutes of Health awards (MH109498 and NS118921) to MG, a National Institutes of Health NIDCD award (5T32DC016903) to EO, and a Sir Henry Dale Fellowship from the Royal Society & Wellcome Trust (101195/Z/13/Z) and a UCL Excellence Fellowship to IHB.

## Methods

### Zebrafish maintenance

Zebrafish were raised and maintained under standard conditions in the BVS Aquatics Facility in the Queen’s Medical Research Institute, University of Edinburgh. Adult and larval animals were maintained on a 14 hours light and 10 hours dark cycle. Embryos were stored in 10mM HEPES-buffered E3 embryo medium or conditioned aquarium water with 0.000001% (w/v) methylene blue at 28.5°C. All experiments were performed under the project license 70/8436 with approval from the UK Home Office. The *myrf*^*UE70*^ line was maintained in a Tupfel Long Fin (TL) wildtype background. Within this manuscript, ‘Tg’ denotes a stable, germline inserted transgenic line.

### Transgenic and mutant lines

The *myrf*^*UE70*^ mutant line was established during this study is described in this manuscript. The following transgenic lines were also used in this study: Tg(mbp:eGFP-CAAX) (21, 63), Tg(mbp:nls-eGFP) (34).

### Generation of *myrf*^*UE70*^ mutants

A freely available guide selection tool (http://crispr.mit.edu) was used to select sgRNA sequences against the second exon of the zebrafish *myrf* gene. sgRNA (target sequence CATTGACACCAGTATCCTGG) was synthesised using DNA template oligomers (5’-AAAGCACCGACTCGGTGCCACTTTTTCAAGTTGATA ACGGACTAGCCTTATTTTAACTTGCTATTTCTAGCT CTAAAACCCAGGATACTGGTGTCAATGCTATAGTGA GTCGTATTACGC-3’) (Integrated DNA Technologies, Belgium) consisting of DNA coding for the T7 promotor, DNA recognition sequence (sgRNA variable region) and the sgRNA scaffold. sgRNA synthesis was performed using Ambion MEGAshortscript T7 Transcription Kit (Thermo Fisher Scientific) and the synthesised DNA oligomers as template. Transcribed sgRNA was purified using Ambion MEGAclear kit (Thermo Fisher Scientific). The expression vector for Cas9 protein, pCS2-nCas9n (Addgene plasmid #47929) (64), was used to transcribe Cas9 mRNA using the mMESSAGE mMachine SP6 kit (Thermo fisher Scientific) and purified using an RNeasy mini kit (Qiagen). Injection solutions were prepared with a final concentration of 300ng/µl nCas9n mRNA and 10ng/µl sgrNA in nuclease free water and 0.05% (w/v) phenol red (Sigma Aldrich). Wildtype embryos were injected at the single or two cell stage with 1.5nL injection solution. Injected F0 animals were raised to adulthood and outcrossed to wildtype animals to create F1 offspring. Clutches of F1 offspring were raised to adulthood and genotyped to identify heterozygous carriers of function disrupting mutant alleles. *myrf*^*UE70*^ refers to a specific allelic mutation consisting of the deletion of two cytosine nucleotides and insertion of a single adenine nucleotide (wildtype sequence: 5’-CCAGTATCCTGGAGGAATA-3’; *myrf*^*UE70*^ mutant allele: 5’-CCAGTATATGGAGGAATA-3’).

### Genotyping

Tissue was genotyped using primers myrf-f (5’-AACTGTGCGTAGGAACACGATA-3’) and myrf-r (5’-TGGACCTCCGTGAAACAACTG-3’) in a standard PCR reaction. The PCR product was digested using restriction enzyme PspGI (New England Biolabs), which cleaves wildtype product into 131bp and 157bp fragments. The mutant product remains uncut as the *myrf*^*UE70*^ allele contains a frameshifting indel which abolishes the PspGI cutting site. PCR products were visualised on a 2% gel following gel electrophoresis. All analyses were performed blinded to genotype.

### Quantitative RT-PCR

Total RNA was extracted from whole brains of adult *myrf*^*UE70*^ wildtype and homozygous siblings using a modified TRIzol™ RNA extraction protocol (TRIzol™ Reagent, Thermo Fisher Scientific). RNA concentration and integrity were assessed using a nanodrop spectrophotometer (NanoDrop Onec, Thermo Fisher Scientific). RNA clean-up was performed if necessary. cDNA synthesis was performed using Accuscript Hi Fidelity First Strand Synthesis kit (Agilent). The amount of RNA entered into the reaction was normalised between samples. Primers mbp-f (5’-ACAGAGACCCCACCACTCTT-3)’ and mbp-r (5’-TCCCAGGCCCAATAGTTCTC-3’) were used to amplify mbp transcripts within a qPCR reaction (Brilliant III Ultra-fast SYBR Green qPCR Master Mix, Agilent). Transcript levels were detected using Roche Light Cycler 96 (Roche Life Science) with the following amplification protocol: preincubation 95° for 180s, two step amplification 40 cycles: 95° for 10s then 60° for 20s, followed by high-resolution melting. Each sample was run in triplicate. House-keeping gene ef1a was used as a reference gene, using primers ef1a-f (5’-TGGTACTTCTCAGGCTGACT-3’) and ef1a-r (5’TGACTCCAACGATCAGCTGT-3’). The delta-delta CT method was used to quantify expression levels. All values were normalised to wildtypes to provide the relative expression of the gene of interest.

### Transmission electron microscopy

Larval tissue was prepared for TEM using the microwave fixation protocol as previously described (34, 65). For adult tissue, adult zebrafish were terminally anaesthetised in tricaine and perfused intracardially with PBS followed by primary fixative solution (4% (w/v) paraformaldehyde, 2.5% (w/v) glutaraldehyde, 0.1M sodium cacodylate) (Sigma Aldrich). Adults were subsequently incubated in fresh primary fixative solution for 24 hours at 4°C. Spinal cords were dissected and processed using the microwave fixation protocol described for larval tissue. TEM images were obtained using a JEOL JEM-1400 Plus Electron Microscope. Image magnification ranged from 11.2-17kx magnification for larval spinal cords, and 1.7kx for adult spinal cord.

### Single cell labelling

Fertilised eggs from *myrf*^*UE70*^ heterozygous adult in-crosses were microinjected between the single and four-cell stage with 10ng/µl plasmid DNA encoding mbp:mCherry-CAAX (63) and 25ng/µl tol2 transposase mRNA in nuclease free water with 0.05% (w/v) phenol red. Animals were screened at 4dpf for mosaically labelled oligodendrocytes and subsequently imaged. Single cells from any level in the dorsal spinal cord were imaged. Images were obtained in 4 and 6dpf larvae.

### Live imaging

Larvae were anaesthetised in 0.6mM tricaine/MS-222 (ethyl 3-aminobenzoate methanesulfonate salt, Sigma Aldrich) in HEPES buffered E3 embryo medium and embedded in 1.3-1.5% (w/v) low melting point agarose (Invitrogen). All fluorescent images were acquired using a Zeiss LSM 880 confocal microscope with a 20x objective (Zeiss Plan-Apochromat 20x dry, NA = 0.8, Carl Zeiss Microscopy). Z-stacks were obtained through the entire single cell, axon or spinal cord according to each experiment. For time course imaging, a single oligodendrocyte was imaged as at 4dpf. Larvae were then extracted from agarose gel, recovered in embryo medium and maintained with daily feeds and water exchange until imaging of the same cell was repeated at 6dpf. For automated imaging of the entire spinal cord and peripheral nervous system, Vertebrate Automated Screening Technology (VAST) was utilised as described previously (66). Briefly, larvae are arrayed into individual wells of a 96-well plate containing MS-222 treated HEPES buffered E3 embryo media. Fish are loaded and oriented for imaging using a Large Particle (LP) Sampler and VAST BioImager system (Union Biometrica Inc) fitted with a 600µm capillary tube. Embryos are automatically loaded into the capillary, positioned and imaged using an AxioCam 506m CCD Camera, a CSU-X1 spinning disk confocal scanner, a 527/54+645/60nm double bandpass emission filter, 1.6x C-Mount adapter, a PIFOC P-725.4CD piezo objective scanner, W-Plan-Apochromat 10x 0.5NA objective and an Axio Examiner D1. Z-stacks covering the depth of the capillary were acquired using a 4µm z-interval, 3×3 binning and 60ms exposure. Images were acquired using brightfield and the appropriate fluorescent channel. Following imaging, larvae were dispensed into a corresponding well of a 96-well collection plate and whole tissue retained for genotyping. Unless otherwise stated, all confocal images presented in this manuscript represent a lateral view of the zebrafish spinal cord, with anterior to the left and posterior to the right, and dorsal/ventral at the top/bottom of the image respectively. Within experiments, images were obtained using similar laser intensity and optical gain settings. All imaging was performed blinded to genotype.

### Transmission electron microscopy

TEM images were tiled using the automated photo merge tool in Adobe Photoshop 2019. The number of ensheathed axons was counted in one hemi-spinal cord section per larva using the cell counter tool in FIJI ImageJ. Axon caliber is defined as the area of the axon within this manuscript. Axonal area was calculated using the freehand line and measure tool in FIJI ImageJ. Calculation of g-ratio was performed by dividing axonal area by the axonal + associated myelin area.

### VAST

Images obtained using VAST were stitched and processed using FIJI ImageJ software (67) and custom macros (66). Semi-automated oligodendrocyte counts were performed on the maximum intensity projection images (66). Cell count values represent all oligodendrocytes in the spinal cord (dorsal and ventral tracts). Morphometric analysis of larval developmental features was performed on brightfield images. Measurements of ocular diameter, body length and swim bladder height were performed using the line and measure tool in FIJI ImageJ (National Institutes of Health).

### Single cell imaging

Confocal z-stack images were airyscan processed using Zen software (Zeiss). Images were opened in FIJI ImageJ. Cells were included for analysis only if all myelin sheaths were distinguishable. Myelin sheath lengths were measured using the freehand line and measure tools. Myelin sheath number was equivalent to the number of measurements performed. Total myelin per cell was calculated as sum of all myelin sheaths lengths per cell. Abnormal sheaths were defined at sheaths with abnormal elongation profiles, incomplete wrapping or myelin blebbing. For time course experiments, net growth or shrinkage of myelin sheaths was calculated as the average myelin sheath length at 6dpf minus the average myelin sheath length at 4dpf. Where possible, all myelin sheaths per cell were measured. In instances where measurement of all myelin sheaths was not possible (due to other cells coming into close proximity), only isolated myelin sheaths were analysed at each time point. The number of retracted sheaths was recorded, and these sheaths were excluded from sheath growth analysis.

### Electrophysiology

Zebrafish were dissected as described previously in Roy & Ali (2013) (23) to access the Mauthner neuron. In short, 6dpf anaesthetised zebrafish were laid on their sides on a Sylgard dish and secured using tungsten pins through their notochords in a dissection solution containing the following (in mM): 134 NaCl, 2.9 KCl, 2.1 CaCl_2_, 1.2 MgCl_2_, 10 HEPES, 10 glucose and 160mg/ml tricaine, adjusted to pH 7.8 with NaOH. Their eyes as well as lower and upper jaws were removed using forceps to expose the ventral surface of the hind-brain, which was secured with an additional tungsten pin. The motor neurons in the anterior spinal cord were exposed as described by Wen et al (2005) (68). A dissecting tungsten pin was used to remove the skin and the muscle overlaying the motor neurons in a single segment. Following the dissection, zebrafish together with their recording chamber were moved to the rig and washed with extracellular solution containing the following (in mM): 134 NaCl, 2.9 KCl, 2.1 CaCl_2_, 1.2 MgCl_2_, 10 HEPES and 10 glucose with 15 µM tubocurarine. The cells were visualised using Olympus microscope capable of DIC using 60X water immersion NA = 1 objective lens and Rolera Bolt Scientific camera with Q-Capture Pro 7 software. The stimulating electrode filled with extracellular solution was then positioned in the anterior spinal cord lightly touching the exposed neurons underneath. Mauthner whole-cell recordings were performed with thick-walled borosilicate glass pipettes pulled to 6–10 MΩ. The internal solution contained the following (in mM): 25 K-gluconate, 15 KCl, 10 HEPES, 5 EGTA, 2 MgCl_2_, 0.4 NaGTP, 2 Na_2_ATP, and 10 Na-phosphocreatine, adjusted to pH 7.4 with KOH. Upon formation of whole-cell patch clamp, 270s – long recording was performed in the current -clamp configuration. Cell resting membrane potential was established as an average of the first 5 seconds of the recording if the cell did not fire during that time. To measure the conduction velocity along the Mauthner axon, the zebrafish were washed with recording solution containing the following (in mM): 134 NaCl, 2.9 KCl, 2.1 CaCl_2_, 1.2 MgCl_2_, 10 HEPES and 10 glucose with addition of (in µM) 50 AP5, 20 strychnine, 100 picrotoxin and 50 CNQX. The antidromic Mauthner action potentials were recorded following the field stimulation by the stimulating electrode connected to DS2A Isolated Voltage Stimulator (Digitimer) in the spinal cord. 30 consecutive action potentials were recoded every 5 seconds using Clampex 10.7 at 100kHz sampling rate and filtered at 2 kHz using Multi-Clamp 700B. At the end of the recording, images of the zebrafish were obtained with 4X objective and stitched using Adobe Photoshop. The resulting image was then transferred to FIJI and the distance between stimulating and the recording electrode was measured. The conduction velocity of action potential was calculated dividing the distance between the electrodes by the latency from the stimulus artefact to the peak of action potential. Action potential latency and half-width were measured using homebuilt MATLAB script. For the analysis of action potential fidelity consecutive trains of 10 stimuli were delivered at 1, 10, 100, 300, 500 and 1000Hz every 30s. Recordings were made at 20kHz sampling rate and filtered at 2 kHz. The number of action potentials were calculated using Clampfit 10.7 software and the action potential success rate was calculated as a number of action potentials fired out of 10.

### Behavioural assay

Analysis of startle behaviour in 5-6 dpf *myrf*^*UE70*^ mutant and wildtype larvae was performed as previously described (39, 46). Briefly, larvae were placed into individual wells of a 6 x 6 custom made acrylic testing plate containing E3 embryo media. A series of 10 acoustic stimuli (40.6 dBu, 1000 Hz, 3ms duration) were delivered to the plate with an interstimulus interval of 20 seconds. Behaviour was recorded using a high-speed camera (Photron Fastcam Mini UX) at 1000frames/s. Analysis of recorded video footage was performed using FLOTE v2.0 tracking software (39). Larvae that responded to less than 70% of the stimuli were excluded from further kinematic analysis. Average behavioural latency was calculated as an average per larva over all behavioural responses. Short latency C-starts (SLC) and long latency C-starts (LLC) were defined by identifying a latency value (16ms) separating the two peaks of the latency bimodal distribution in wildtype *myrf*^*UE70*^ larvae. Behavioural latency, c-bend duration, initial turn angle, and angular velocity for SLC and LLC events were defined and analysed as previously described (39).

### Experimental design and statistical analysis

Unless stated otherwise, all experiments were performed on 6dpf larvae from adult heterozygous in-crosses. All subjects were the offspring of third generation, or younger, adults. The experimenter was blinded to the genotype of the larvae during all experiments and analysis. The sex of the animals was unknown as sex specification has not occurred at this stage of larval development. All graphs and statistical testing were performed using GraphPad Prism. All data was assessed for Gaussian distribution using a D’agostino Pearson omnibus normality test. Parametric continuous data was analysed using a two-tailed unpaired student’s t test, or two-way ANOVA, according to the number of variables being compared. Non-parametric continuous data was analysed using a Mann-Whitney test. If the number of values were too small to assess for normality, it was assumed that data was non-parametric. Results were considered statistically significant when p < 0.05. Within figures, p values are denoted as follows: non-significant i.e. p> 0.05 ‘ns’, p <0.05 ‘*’, p <0.01 ‘**’, p<0.001 ‘***’, p < 0.0001 ‘****’. Unless otherwise stated, all data was averaged per biological replicate (N represents number of larvae). Throughout the figures, error bars represent mean ± standard deviation for parametric data, and median and interquartile range for non-parametric data. Details of statistical tests, precise p and n values for each experiment are provided in the appropriate figure legends.

## Code accessibility

Custom written code to perform automated cell counts is available in a previous publication (66).

